# Naïve T lymphocytes chemotax to CCL21 but not to S1P-rich serum

**DOI:** 10.1101/2023.02.03.526993

**Authors:** Nicolas Garcia-Seyda, Solene Song, Luc David-Broglio, Christoph Matti, Marc Artinger, Martine Biarnes-Pelicot, Marie-Pierre Valignat, Daniel F. Legler, Marc Bajénoff, Olivier Theodoly

## Abstract

Naïve T lymphocytes traffic through the organism in their search for antigen, alternating between blood and secondary lymphoid organs. Lymphocyte homing to lymph nodes relies on the chemokine CCL21, while exit into efferent lymphatics relies on the sphingolipid S1P. Surprisingly, while both molecules are claimed chemotactic, a quantitative analysis of naïve T lymphocyte migration along defined gradients is missing. Here, we used a reductionist *in vitro* approach to study the real-time, single-cell response of naïve T lymphocytes to CCL21 and S1P-rich serum. Using high-throughput microfluidic and optical micropatterning ad hoc tools, we show that CCL21 triggers long-range chemotaxis whereas S1P-rich serum does not. Instead, S1P-rich serum triggers a transient polarization that may represent a brief transmigration step through exit portals. Our data thus validate naïve T lymphocyte chemotaxis towards CCL21 but not S1P, which complements *in vivo* observations and is of interest for a better tailoring of immunosuppressive drugs.

## Introduction

Chemokinesis: random migration triggered by a soluble cue.
Haptokinesis: random migration triggered by an adsorbed cue.
Chemotaxis: directed migration along a soluble cue shaped as a gradient.
Haptotaxis: directed migration along an adsorbed cue shaped as a gradient.

Naïve T lymphocytes circulate through the organism in the search for cognate antigens, thereby alternating between secondary lymphoid organs (SLOs), lymphatics and the blood^1^. Entry to and homing within SLOs are dependent on the CCR7 receptor, present on lymphocytes, and its cognate ligand CCL21 produced by stroma cells in lymphoid tissues^1,2^. Egress from SLOs is in turn dependent on the S1PR1 receptor and its cognate ligand S1P, a small sphingolipid abundant in blood and lymph^3^. The lymphocyte transit time through lymph nodes is thus controlled by a balance between CCL21-controlled recruitment and retention, and S1P exit signals^4^. For this reason, disruption of the S1PR1-S1P signaling axis represents an immunosuppressing therapy applicable to a wide range of pathologies, including multiple sclerosis, transplant rejection, diabetes and cancer^5,6^. Surprisingly, while there is a general consensus that both CCL21 and S1P carry their functions through chemotaxis, a quantitative analysis of naïve T lymphocyte migration along defined CCL21 and S1P gradients is missing.

In the case of CCL21, lymph node gradients have been reported across organ peripheries: Across B cell follicles^7^, along interfollicular regions^8^, and along medullary cords^9^. All three increase in concentration towards the central parenchyma, suggesting a single and common source: the T cell zone. Due to its positively charged C-terminal tail, CCL21 interacts with and is retained by extracellular matrix (ECM) components such as heparan sulfate or collagen^10–13^. This capacity prevents it from being washed away during immunohistological sample preparation, allowing its visualization as a gradient, and it has been claimed that only such immobilized CCL21 triggers naïve T lymphocyte migration^14^. *In vivo*, chemotaxis of naïve T lymphocytes has only been recorded along medullary cords^9^. Other reports suggest instead random migration, at least in other regions of the lymph node^15–18^. However, a drawback of *in vivo* experiments is that they cannot prove whether chemotaxis is triggered by a single visualized gradient or by additional yet unspecified gradients. For instance, CCL19 is another homeostatic CCR7 ligand present in lymph nodes and triggering naïve T lymphocyte chemotaxis *in vitro^19^*, but its distribution *in vivo* remains unknown because it does not bear a ‘sticky’ tail and does not become immobilized^20^. Since T cell zone stromal cells simultaneously produce CCL19 and CCL21^21,22^, both chemokines are necessarily overlapping, making it difficult to judge which one is guiding naïve T lymphocytes *in vivo*. Also, the fact that a chemokine may be simultaneously immobilized or soluble, coupled to the impossibility to reveal the soluble pool by staining, may further hinder the deciphering of traffic mechanisms. Such co-existences of immobilized and soluble chemokine pools have been demonstrated for the B cell zone chemokine CXCL13^23,24^. In the case of CCL21, Dendritic Cells (DCs) cleave its C-terminal tail transforming it into a soluble pool that triggers chemotaxis of these cells^25^, but has additional unique features as compared to the other CCR7 ligands^26^. In this context, a model of chemokine cloud was tentatively proposed in which molecules are trapped as local ‘soluble depots’ within the glycocalix^27^. Regarding naïve T lymphocytes, it remains unclear to what extent CCL19 and CCL21 guide them *in vivo*, and whether CCL21 does it as a soluble or immobilized pool.

For S1P, the perspective is even more complex due to its pleiotropic functions, and since its soluble and lipidic nature impairs molecular labeling and gradient visualization. An elegant tool was recently developed in which S1P presence is deduced from the internalization rate of its receptor, which allowed for gradient identification in the spleen^28^. However, while the same authors reported higher S1P concentrations in lymph node medullary cords than in the T cell zone, they failed to detect a gradient within or between those two areas^29^. In addition to the lack of *in vivo* gradient identification, the percentage of naïve T lymphocytes responding *in vitro* to S1P is strikingly low, typically below 10% ^4,30–34^. While it was claimed a consequence of its receptor’s fast desensitization, this number was recently increased to almost 20% when lymphatic endothelial cells (LECs) were added to the experiments^35^, consistent with those cells controlling a transmigration step towards S1P, rather than long-distance chemotaxis towards it. In the same line, other functions have been suggested for S1P such as migration inhibition alone^36^ or migration inhibition with modulation of adhesion^37^. Another report defends a stromal gate model where S1P acts mainly on LECs, to allow or block lymphocyte transmigration without otherwise affecting their migration^38^. Finally, an *in vivo* report revealed that naïve lymphocytes randomly approach cortical sinus exit points, with no apparent chemotaxis involved^16^. Altogether, while chemotaxis to S1P is the prevailing model for naïve T lymphocyte exit from lymph nodes, the standing evidence is conflicting.

Chemotaxis towards CCL21 and S1P remains to be faithfully demonstrated with a reductionist *in vitro* experiment where cells would migrate along a single, controlled gradient. However, the typical off-the-shelf tools in biology or immunology labs, the Transwell assays, do not properly and selectively probe chemotaxis. Transwell assays consist of two overlaid chambers, the top one containing cells and the bottom one the molecule being tested. The chambers are separated by a porous membrane through which cell transmigration is scored. While easy to handle, these assays offer no control over the gradient shape nor the moment of its arrival (time zero). They are an endpoint assay with no mechanistic insight due to the lack of *live* imaging, score transmigration through a 10-50 μm thick porous membrane without information on longer-distance gradients, and in the absence of proper controls are unable to distinguish effects of transmigration, chemokinesis or chemotaxis^39^. Moreover, in the case of S1P such controls are uninformative due to the fast receptor desensitization, which precludes coincubating the molecule with the cells in the upper chamber.

The uncertainty on CCL21 and S1P guiding properties would be finally solved with *live* imaging, accessible with microfluidic tools. However, microfluidic devices for gradient generation are generally based on flow or do not control residual drifts, therefore washing away weakly or non-adhesive cells. This is the case for naïve T lymphocytes, claimed to be non-adhesive on ICAM-1 unless subjected to mild shear stress^14^. To circumvent this caveat, we recently developed a microfluidic device for gradient generation in the absence of flow and used it to prove naïve T lymphocyte chemotaxis towards CCL19^19^. Here, we used high-throughput microfluidic and protein printing ad hoc approaches to dissect the response of naïve T lymphocytes to immobilized and soluble CCL21 and S1P-rich serum, at the single cell level. We first show that both adsorbed and soluble CCL21 trigger naïve T lymphocyte hapto- and chemo-kinesis, respectively, when presented as homogeneous chemokine fields. Next, we show that naïve T lymphocytes do adhere on ICAM-1 substrates, though in a weak and intermittent way which is not stabilized by shear stress. We finally demonstrate that CCL21 gradients trigger naïve T lymphocyte hapto- and chemo-taxis, while under similar conditions S1P-rich serum does not. Serum triggers instead a transient polarization which is consistent with a short transmigration step, rather than a long-distance attraction.

Importantly, there is a long-standing call for better understanding the human immunology^40,41^. With most of the above-mentioned evidence arising from mouse models and a recent report indicating differences between both species^42^, plus the numerous therapeutic opportunities expected from a better understanding of the S1PR1-S1P signaling axis, we carry our studies on naïve T lymphocytes purified from healthy human donors.

## Results

### CCL21 triggers naïve T lymphocyte hapto- and chemo-kinesis

Based on *in vitro* live imaging, it has been claimed that CCL21 does not stimulate naïve T lymphocytes unless adsorbed on a substrate^14^. Since our microfluidic device creates soluble gradients based on diffusion, we first tested whether CCL21 triggered migration of human naïve CD4^+^ T lymphocytes while in bulk solution only. We analyzed the behavior of cells in non-adherent single channels either coated with the chemokine, rinsed and blocked with BSA (Fig 1A, adsorbed CCL21), or coated with pluronic F127, which keeps the chemokine and cells in solution^43^ (Fig 1B, bulk CCL21). We observed cell polarization and random migration in both conditions, demonstrating the molecule’s hapto- and chemo-kinetic potential (Fig 1 and Movie 1). To verify that CCL21 had not permanently adsorbed (immobilized) on the F127 substrate, the channel was rinsed at the end of the experiment and fresh cells were added. We observed only 2% migrating cells, proving that the chemokine had been rinsed and thus suggesting that the observed effect was triggered from molecules in the bulk solution. Based on these results, we conclude that CCL21 triggers efficient cell migration when presented in solution, and therefore can be used in our microfluidic device for studying naïve T lymphocyte chemotaxis. In addition, migration in the absence of adhesion reveals the swimming capacity of naïve T lymphocytes, as previously reported for effector T lymphocytes, amoeba and neutrophils^43–45^.

**Fig 1.**
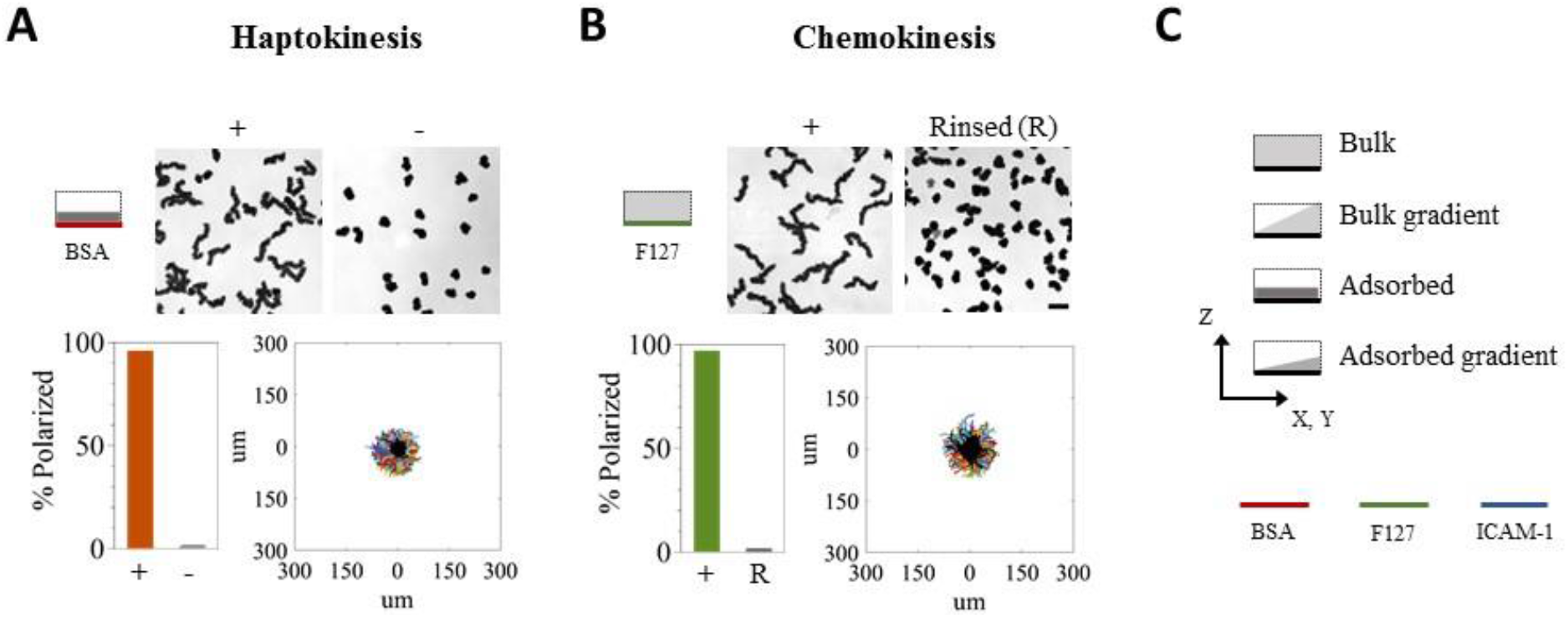
CCL21 triggers hapto- and chemo-kinesis of naïve T lymphocytes. Top row, bright field 10-minute time projections of cells in the presence (+) or absence (-) of CCL21 adsorbed on BSA (A, haptokinesis) or kept in bulk solution over an antifouling F-127 substrate (B, chemokinesis). Bottom, Quantification of cell polarization in each condition and trajectories aligned in the origin for cells tracked over a 16-minutes period, in the presence (colored lines, > 500 tracks are shown) or absence (overlaid in black, > 500 tracks are shown) of chemokine. For the last condition, bulk CCL21 was rinsed (R) and fresh cells were added to detect potentially adsorbed chemokine. Data is from one representative donor, nBSA = 2, nF127 = 3 donors. Scalebar = 30μm. (C) Cartoon description as used in all subsequent figures, describing stimulus nature and substrate color-code. The dotted frames represent the limits of the experimental chamber, from a transversal viewing point.

### Naïve T lymphocytes intermittently adhere on ICAM-1

It has also been claimed that naïve lymphocytes do not adhere on ICAM-1 in the absence of shear stress^14^. However, when we imaged cells in single channels coated with ICAM-1 and CCL21 we noted that they explored a greater surface area (Fig 2A and Movie 2). Instant speed analysis revealed two populations on ICAM-1 substrates: one of low speed, equal to that of cells in the absence of adhesion, and one of higher speed, likely corresponding to cells adhering on the ICAM-1 (Fig 2B). We therefore performed Interference Reflection Microscopy (IRM) to characterize such populations. With this imaging technique, cells in close contact to the substrate display destructive optical interference leading to an intensity darker than the background, whereas non-adherent cells present constructive optical interference leading to a brighter intensity. While cells migrating on BSA barely presented adhesion fingerprints, cells on ICAM-1 sequentially alternated between adherent and non-adherent states (Fig 2C). Single cell analysis of instant speed and adhesion area further revealed that temporal adhesion is indeed a prerequisite for fast migration (Fig 2C). Intermittent adhesion did not arise from a shortage of adsorbed chemokine since similar results were obtained when the chemokine was kept in bulk solution, at high concentration, over the ICAM-1 substrate (Fig 2D and Movie 3). Finally, application of a 0.2 dyn shear stress did not stabilize adhesion^14^ but rather washed cells away, likely when switching into the swimming regime (Fig 2E and Movies 4 and 5). Based on these results, we conclude that naïve T lymphocytes do adhere on ICAM-1, though in an intermittent fashion which is not stabilized by shear stress nor chemokine availability.

**Fig 2.**
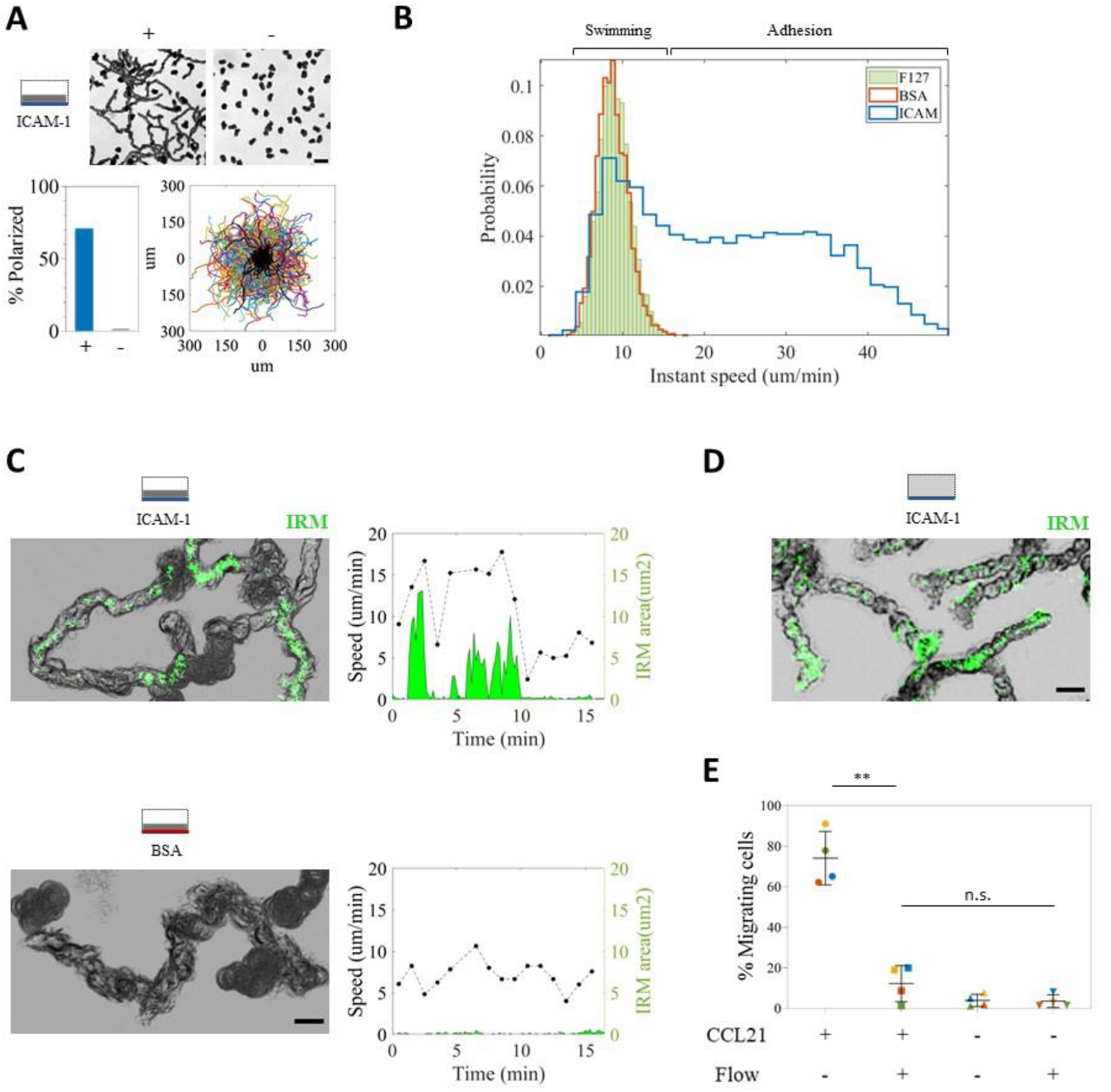
Naïve T lymphocytes intermittently adhere to immobilized ICAM-1. (A) Top row, bright field 5-minute time projection of cells in the presence (+) or absence (-) of CCL21 adsorbed on ICAM-1. Scalebar = 30μm. Bottom, Quantification of cell polarization in each condition plus trajectories aligned in the origin for cells tracked over a 16-minute period, in the presence (colored lines, >1000 tracks) or absence (overlaid in black, >1000 tracks) of chemokine. Data is from the same donor as in Fig 1, nICAM-1 > 3 donors tested. (B) Instant speed distribution calculated over 1-minute intervals, for migrating cells in Fig 1 and 2A. n = 11709, 9256 and 5095 values calculated for ICAM-1, BSA and F127, respectively. (C) Left, overlaid bright field (grey) and inverted IRM (green = adhesion patch) time projections, to reveal adhesion fingerprints while migrating on the indicated substrates. nICAM-1 > 3 donors tested. Scalebar = 10μm. Right, single cell analysis of instant speed (over 1-minute intervals) and adhesion area for a representative cell on each substrate. (D) Overlaid bright field (grey) and inverted IRM (green = adhesion patch) time projections for cells migrating on an ICAM-1 substrate with 1μg/ml CCL21 kept in bulk solution. Scalebar =10μm. n > 3 independently tested donors. (E) Percent of migrating cells remaining on ICAM-1 substrates with or without adsorbed CCL21 after addition of 0.2dyn flow. Each color represents an independently tested donor. ** indicates a p-value < 0.01 in a multiple comparison ANOVA test.

### CCL21 gradients trigger naïve T lymphocyte hapto- and chemo-taxis

Given that bulk CCL21 triggered naïve T lymphocyte chemokinesis, we next tested whether it also triggered their chemotaxis by using our microfluidic device for soluble gradient generation^19^. In our setup, cells are injected in a central channel and their behavior is recorded in response to a gradient established by diffusion (Fig 3A). Parallel channels on each side are used as the chemoattractant source and sink; they are separated from the central one by a double array of trapezoidal pillars holding permeable agarose barriers, which allow chemokine diffusion while dampening flow across them. A mild flow assures chemokine replenishment and removal at the source and sink channels, respectively, while the width of the central channel imposes the gradient steepness, with profiles spanning 500 to 1000 um in length^19^. When CCL21 was applied as a soluble gradient, we observed strong directional migration towards increasing chemokine concentrations (Fig 3B and Movie 6). However, because the chemokine adsorbs on various substrates (Fig 1 and 2), this effect could arise from a combination of the applied soluble gradient plus a potential haptotactic one building up during the experiment, due to molecules instantly captured on the substrate. We thus turned into self-made chemokine versions^46^ to identify each contribution.

**Fig 3.**
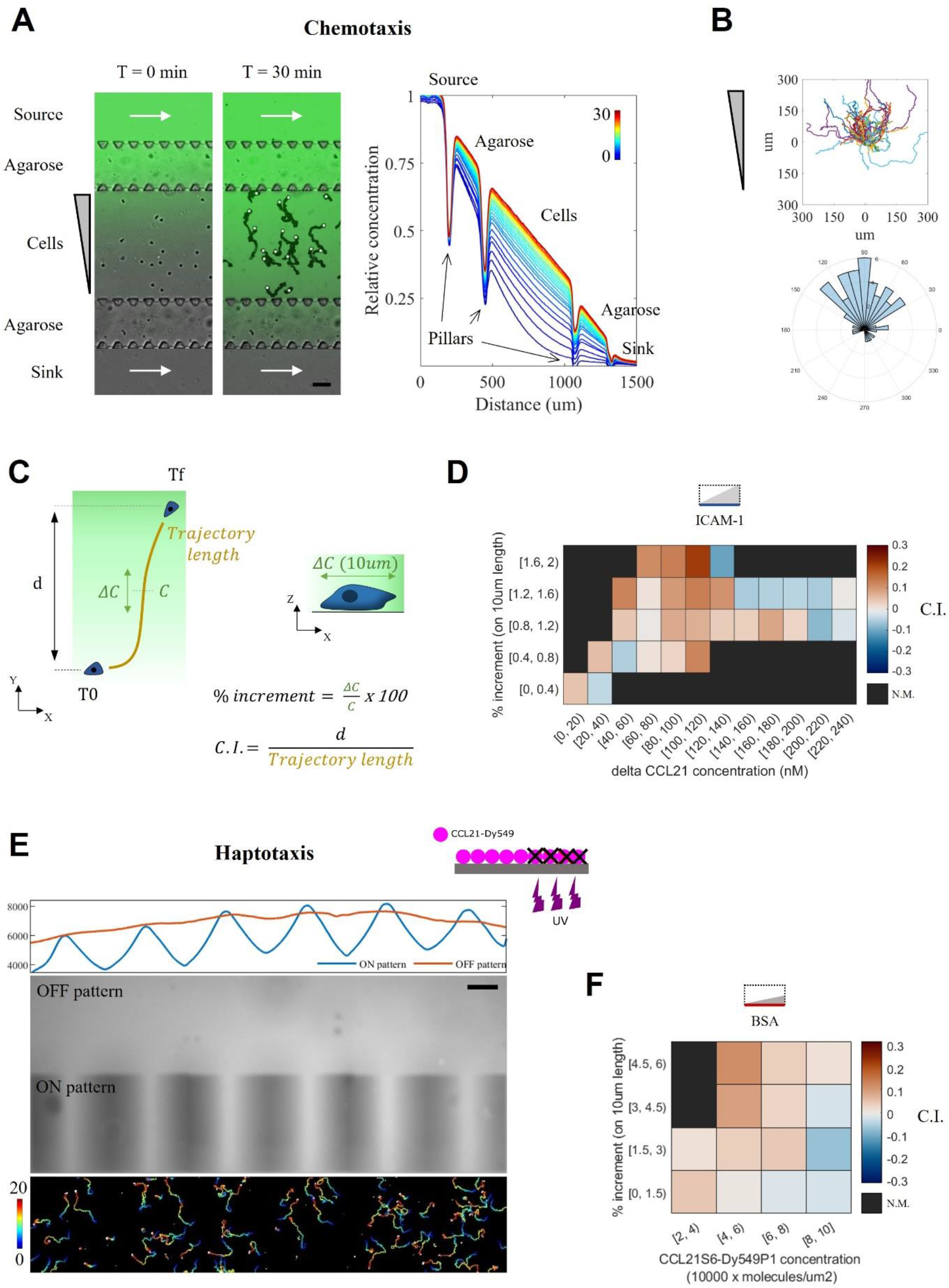
CCL21 gradients trigger hapto- and chemo-taxis of naïve T lymphocytes. (A) Left, overlaid bright field and fluorescent signal for an illustrative CCL21 gradient at time zero and after 30 minutes of acquisition. For the later, bright field images were projected to highlight cell trajectories, the white circles indicating their final position. White arrows indicate flow direction in the side channels, replenishing the source and clearing the sink to keep them at maximum and minimum chemokine concentrations, respectively. Agarose barriers, held by trapezoidal PDMS pillars, allow chemokine diffusion while hampering fluid flow across the central channel. Fluorescent 10 Kda Dextran is used to verify gradient establishment and lack of flow. Scalebar = 100μm. Right, normalized gradient profiles taken at 1-minute intervals, the color-code representing time from 0 to 30 minutes. Opaque PDMS pillars appear as fluorescent signal drops. Cells experience increasing concentrations of chemokine, ranging from 0 to 75% of the concentration at the source. (B) Trajectories aligned in the origin and angle histogram for a total of 55 cells from one representative donor, tracked over 50 minutes. N = 4 donors tested. (C) Cartoon illustrating the calculation of the chemotactic index (C. I.) as a ratio between the net displacement along the gradient direction (d) and the trajectory length. Each value is then tagged with the local chemokine concentration (C) and slope (ΔC) over 10μm length, the typical body size for a naïve T lymphocyte. (D) Heatmap for the chemotactic Index (C.I.) as a function of ΔCCL21 bulk concentration and gradient slope for 3 independently tested donors (n = 215 tracks, 6422 C.I. values calculated over 1-minute intervals). N.M. = Not Measured or below a threshold of 25 values. (E) Top, cartoon illustrating the subtractive printing protocol. Below, fluorescent image and profiles along the patterned area (ON pattern) or out of it (OFF pattern). Bottom, cell trajectories on the patterned area, color-coded with time over a 20-minute period. (F) Heatmap for the chemotactic Index (C.I.) as a function of chemokine substrate concentration and gradient slope for 3 independently tested donors (n = 6809 tracks, 7878 C.I. values calculated over 1-minute intervals). N.M. = Not Measured or below a threshold of 25 values.

To determine the soluble contribution we used a truncated variant, CCL21^24-102^ or ΔCCL21, lacking the C-terminal basic motif and therefore expected to remain exclusively in bulk solution. Because its diffusion is represented by a similar weight FITC-Dextran tracer, fine analysis of Chemotactic Index (C.I.) versus gradient concentration and slope was achieved (Fig 3C). Chemotaxis was maximum at high slopes and 80-120 nM concentrations, but detectable from a 0.4% increment over a cell body-length (Fig 3D). To determine the haptotactic contribution, we then used a fluorescently labelled CCL21, CCL21-S6^Dy549P1^. Starting from a homogeneously adsorbed chemokine substrate as in Fig. 1A, we used a subtractive printing protocol^47,48^ to degrade chemokine functionality and create patterns of interest in various slopes and intensity ranges (Fig 3E). Since leukocyte migration is biased by gradients of adhesion ligands^47^, we performed these experiments in the absence of adhesion. The fluorescent signal was then transformed into number of adsorbed molecules with the aid of a calibration curve (Supplementary fig. 1). When correlated to the calculated C.I. values, haptotaxis was identified at high slopes and 4-6 x 10^4^ molecules/um^2^ (Fig. 3F). Based on these results, we conclude that CCL21 effectively triggers naïve T lymphocyte hapto- and chemo-taxis.

### Serum modulates S1PR1 surface expression on naïve T lymphocytes

Naïve T lymphocytes are assumed to exit lymph nodes following a gradient of S1P. However, directed migration towards S1P was never imaged, neither *in vivo* nor *in vitro*. We therefore decided to challenge this idea using our microfluidic device. Due to its lipidic nature, S1P is carried in blood by albumin and apolipoprotein M, each of them having apparent distinct functions and cellular targets^49,50^. Those carriers are not yet elucidated for lymph, thus it is not known in which state naïve T lymphocytes encounter and sense S1P at cortical sinus exit points, which might explain the low transmigration values reported in the literature. Following this consideration, and since lymph is mixed with blood at the thoracic duct and both fluids trigger similar S1PR1 desensitization^51^, we tested autologous donor serum as the source of bioactive S1P. In this way, we also sought to reduce inter-donor variability due to the use of human cells. Because experiments in mice proved that S1PR1 desensitizes within minutes of exposure^30^, we first sought to characterize its dynamics in human cells. In agreement with the literature^52^, naïve T lymphocytes did not express S1PR1 when directly stained in blood, though resensitization occurred during cell purification (Fig 4A). When incubated in the absence of fetal calf serum, cells remained viable (Supplementary fig. 2A) and S1PR1 expression reached a plateau with 84 ± 8% resensitized cells (Supplementary fig. 2B). When resensitized cells were then exposed to autologous human serum, we observed a fast, concentration dependent S1PR1 desensitization within 5 minutes (Fig 4B). These results thus validated serum as a source of bioactive S1P, which was used for subsequent experiments.

**Fig 4.**
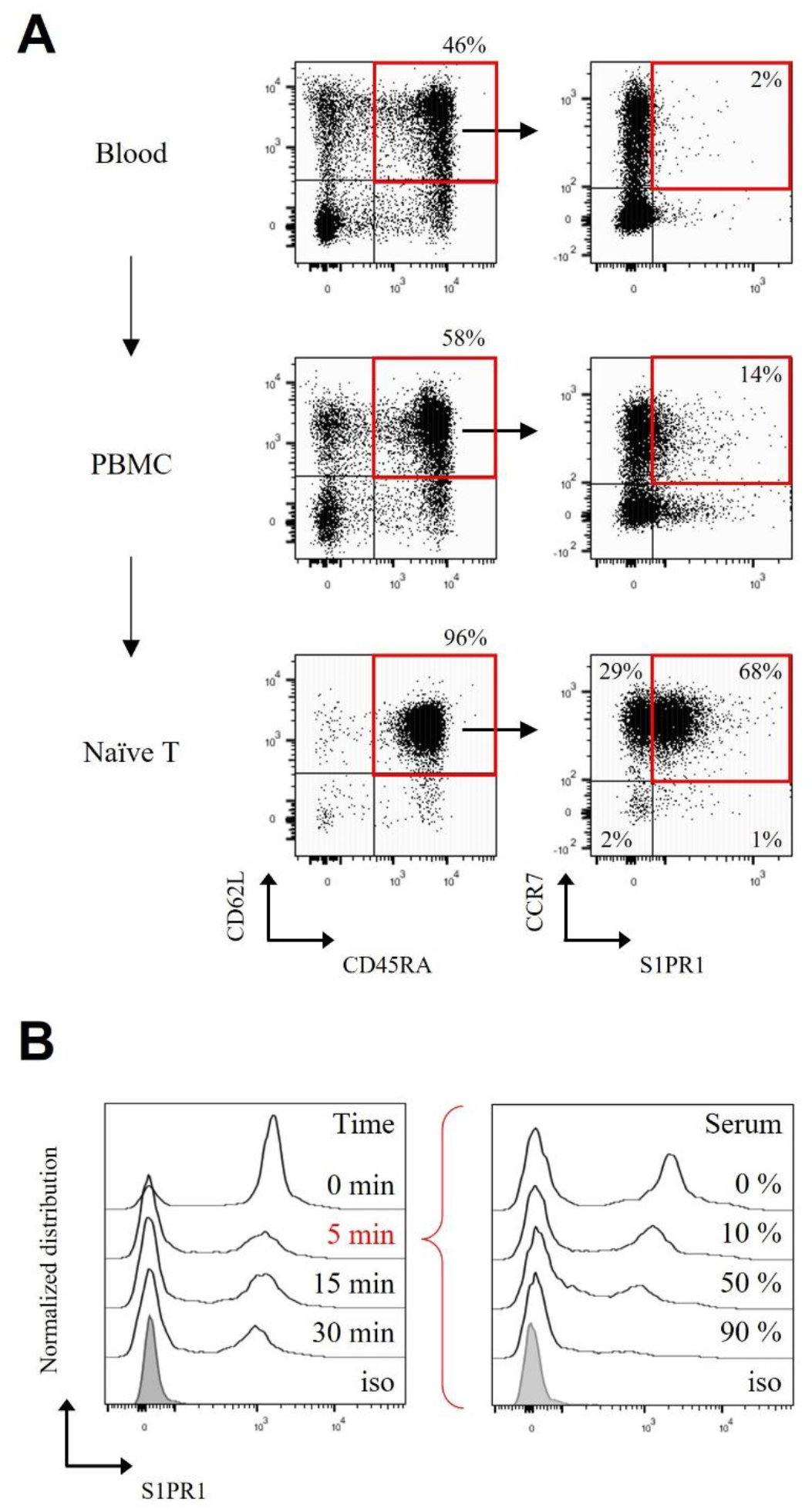
Serum modulates S1PR1 surface expression on naïve T lymphocytes. (A) S1PR1 live staining performed on whole blood cells, PBMC’s and naïve T lymphocytes, along the purification process. Arrows indicate the gating strategy. (B) Left, S1PR1 modulation on cells exposed to 50% serum concentration and then fixed at the indicated timepoints. Right, S1PR1 modulation on cells exposed 5 minutes to the indicated serum concentrations and then fixed. Gray histograms correspond to isotype controls. Data representative from 1 out of 3 and 2 independently tested donors, respectively.

### Serum triggers transient polarization and chemokinesis of a fraction of cells, but not chemotaxis

We first analyzed the response of resensitized cells to serum in single channels coated with ICAM-1. We observed a mild but significant effect for 10% serum concentration, with 12 ± 7% polarizing cells (Fig 5A). IRM imaging proved migrating cells were intermittently adhering, ruling out the hypothesis of adhesion inhibition (Fig 5B and Movie 7). In addition, when cells were simultaneously exposed to both S1P-containing serum and CCL21, no apparent inhibition of cell migration nor adhesion was observed (Supplementary fig 3). In line with a faster receptor desensitization at high serum concentrations (Fig 4B), we observed only 3.5 ± 1% polarized cells with 90% serum concentration. Because many of the responding cells were already polarized when starting the acquisition, we assume the response occurred and was completed during cell sedimentation, typically few minutes long. This result highlights the importance of controlling and visualizing the instant of stimulus addition (time-zero), achievable only in controlled microfluidic setups.

**Fig 5.**
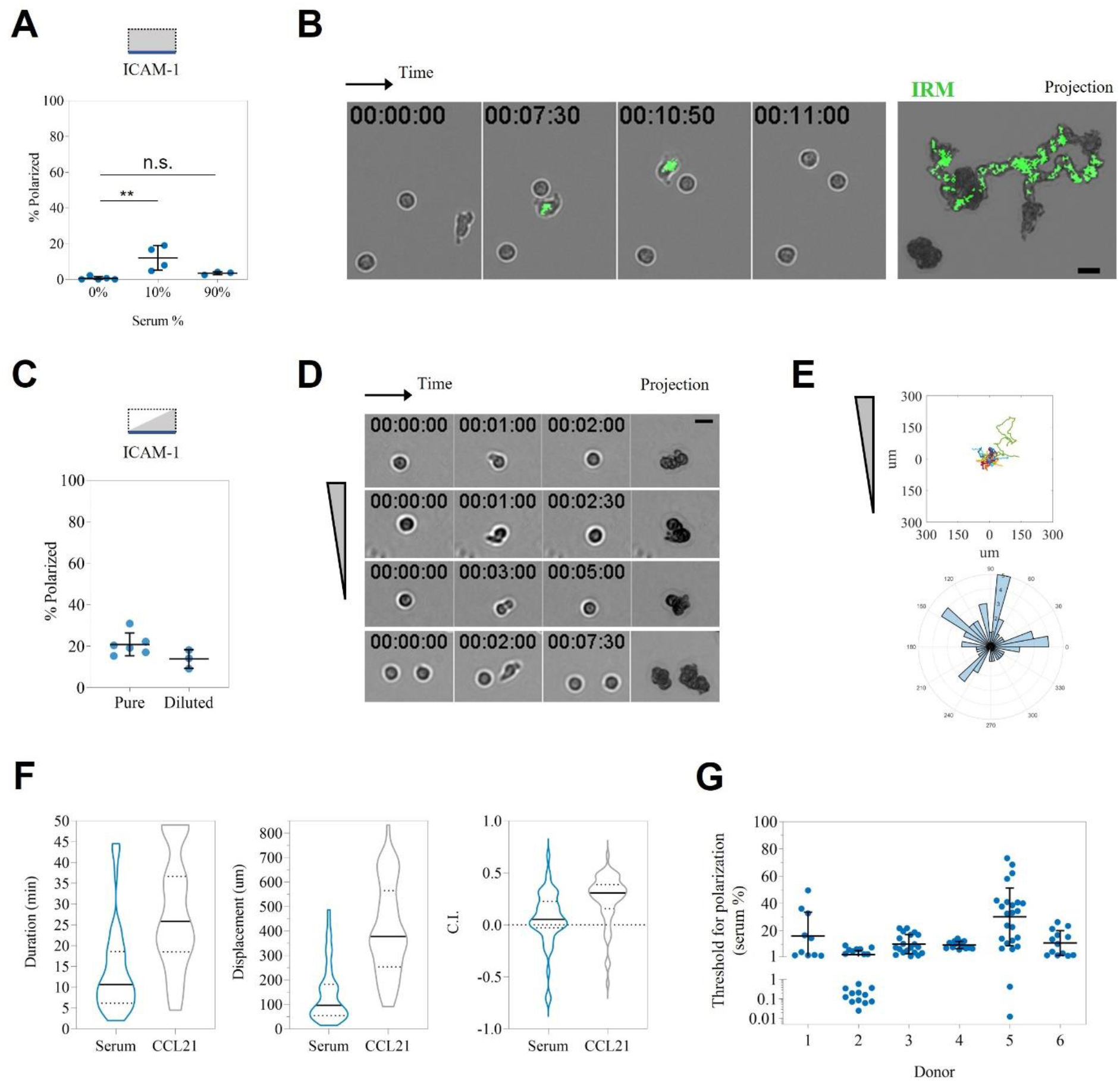
Serum triggers transient polarization and chemokinesis of a fraction of cells, but not chemotaxis. (A) Cell polarization in single channels in the presence of the indicated serum concentrations. A total of 664 cells from 5 independently tested donors were imaged. (B) Time sequence and overlaid projections for bright field and inverted IRM imaging (green = adhesion patch) for a group of resensitized cells in 10% serum. (C) Quantification of cell polarization upon serum gradient arrival. 620 cells from 6 independently tested donors were exposed to pure serum (0 to 70% concentration range), while 395 cells from 3 independently tested donors were exposed to diluted serum (0 to 15% concentration range). (D) Time sequence and projections of 4 representative cells shortly polarizing upon serum arrival, but without net displacement. (E) Trajectories aligned in the origin and angle histogram for the cumulated 49 cells amenable for tracking. (F) From left to right, distribution of duration, displacement, and chemotactic index (C.I.) for those 49 tracks. As a comparison, same parameters for the 55 tracks shown in Fig 3B (CCL21 gradient, 1 representative donor) are plotted. (G) Threshold for cell polarization, defined as the serum concentration at the instant of symmetry breaking, for the 6 donors exposed to undiluted serum gradients. Values below 1% are plotted in a log scale. Each point in the dot blots represents one independently tested donor, except in (G) where they represent one cell. Scalebars = 10μm. ** indicate a p-value < 0.01 measured by multiple comparison ANOVA test.

We then exposed resensitized cells to controlled serum gradients in our microfluidic device. By visualizing the moment of serum arrival, we observed a slightly stronger effect than in single channels, with 21 ± 6 % cells polarizing upon serum arrival (Fig 5C). For many of them though the effect lasted few minutes, shortly polarizing on the spot without a net displacement (Fig 5D and Movie 8). Indeed, only 48 out of 620 imaged cells (7.7 %) displaced more than 20 μm (2 body lengths) and were tracked (Fig 5E). Track duration was short, with a median of 11 minutes, therefore limiting cell displacement to 100μm (Fig 5F and Movie 8). As a comparison, CCL21 tracks ended either when the cells reached the channel’s upper limits or when the movie, typically 50 minutes long, was over. Migrating cells did not exhibit a marked directionality towards the serum source, albeit the distribution of C.I. values was slightly skewed towards it. Because our gradients are built by gradual arrival of diffusing compounds, a threshold for polarization was determined by measuring serum concentration at the time of symmetry breaking. We observed a strong variation between and within individual donors, however many cells proved sensitive to less than 10% serum concentration (Fig 5G). Since receptor saturation at high concentrations lowers the Chemotactic Index (as exemplified with ΔCCL21 in Fig 3D) and in the case of serum causes faster desensitization (Fig 4B) with an expected lower number of responding cells, we finally diluted the serum in culture medium and exposed cells to unsaturating gradients. The observed effect though was weaker, with only 14 ± 5 % polarizing cells (Fig 5C). Altogether, we conclude that under the same experimental conditions in which CCL21 triggers long-range chemotaxis of most cells, S1P-rich serum does not. Instead, serum is shortly chemokinetic on a small fraction of cells, while the transient polarization and lack of displacement of the remaining fraction of cells rather points towards a ‘decision making’ function.

### Instant stimulus arrival confirms naïve T lymphocyte transient polarization to serum

Gradient experiments attained a maximum of only 21% responding cells (Fig 5C), as opposed to the global S1PR1 desensitization observed in flow cytometry (Fig 4D). Since in our device serum is gradually arriving by diffusion, we hypothesized that non-reacting cells may desensitize S1PR1 before reaching the threshold for a response. We therefore sought to instantly expose cells to serum as in Fig 5A, but with time-zero visualization and control. To achieve this, cells were captured on the substrate to prevent their flushing upon serum addition, in an ‘open-well’ setup to allow a fast stimulus arrival while recording cell behavior (Fig 6A). We used our optical micropatterning tool to generate an array of 240 circular capture spots coated with α-CD45RA antibodies, to increase the experimental throughput while keeping a single-cell analysis and performed quantitative morphometry of cell contours to detect changes in their polarization state. When cells were first exposed to CCL21, we observed a fast and stable polarization of 71 ± 7% cells, validating the method (Fig 6B and C). When the equivalent experiment was performed with serum, polarization was fast but transient, with 59 ± 5% and 54 ± 14% polarized cells for 10% and 100% serum respectively, and no statistical difference as compared to CCL21 control (Fig 6B, C and Movie 9). Based on these data we conclude that fast exposure to serum transiently polarizes naïve T lymphocytes, providing a qualitative signal consistent with a ‘decision making’ function which is substantially different from the durable guiding signal triggered by CCL21 (Fig 7).

**Fig 6.**
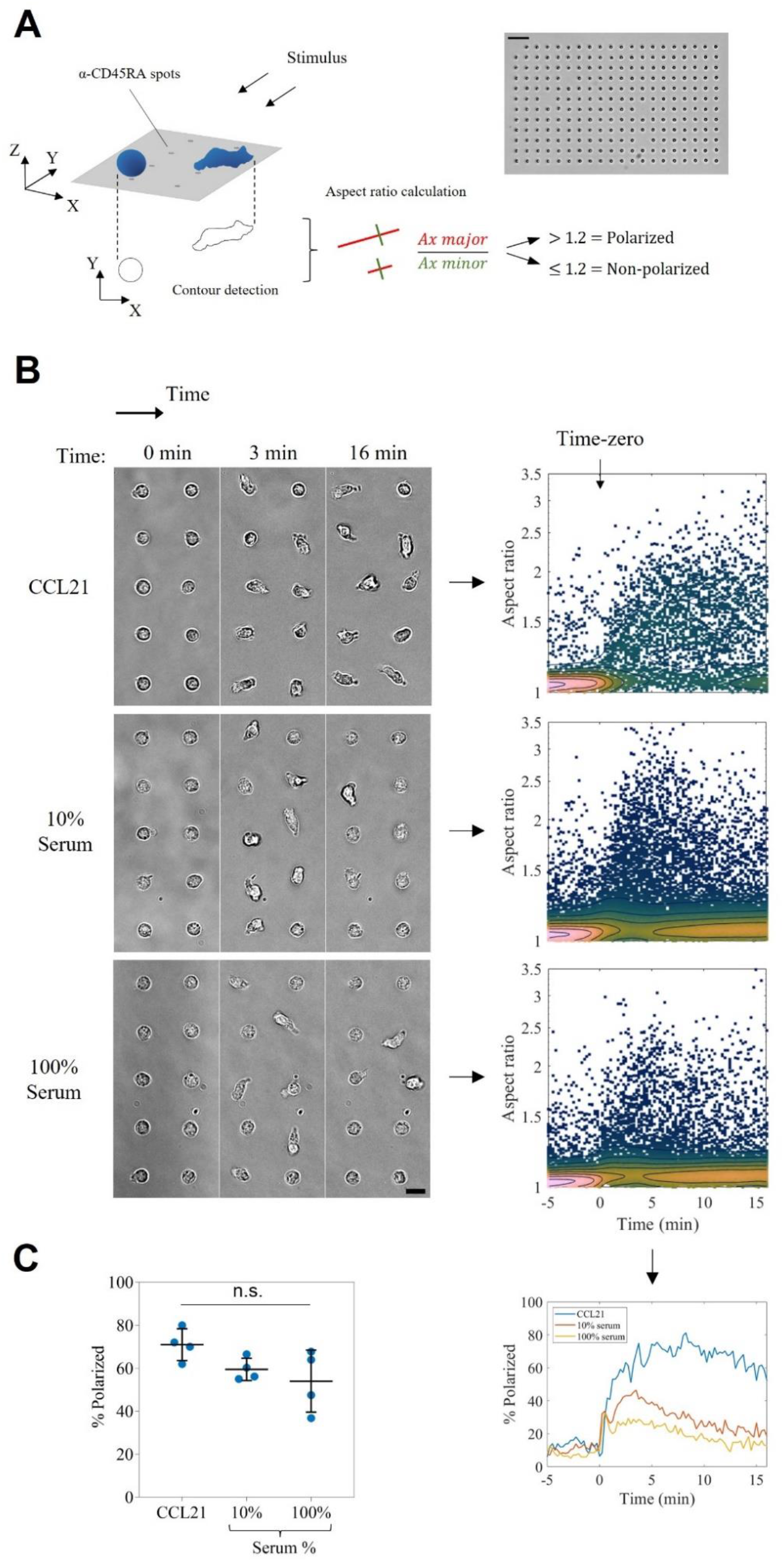
Instant stimulus arrival confirms naïve T lymphocyte transient polarization to serum. (A) Left side, cartoon exemplifying the ‘open-well’ experimental setup and the morphometric analysis used to score individual cell polarization states. Cells were considered polarized when their aspect ratio was higher than 1.2. Right side, low magnification bright-field image of a substrate after cell capture. Scalebar = 50 μm. (B) Left side, high magnification bright-field images of cells from one representative donor at Time-zero, 3 and 16 minutes after the indicated stimulus arrival. Scalebar = 10 μm. Right side, density scatter plots of cell aspect ratio as a function of time for each condition tested. Each dot corresponds to a single cell-contour and is colored according to the local density of events. Below, cell polarization as a function of time for the three datasets. Time-zero corresponds to the instant of stimulus addition. (C) Quantification of cell polarization during the whole acquisition period, values are higher than in (B) due to asynchronous cell response. Each condition was tested on 4 donors, n = 447, 652 and 440 imaged cells for CCL21, 10% and 100% serum, respectively. n.s. = not significant (multiple comparison ANOVA test).

**Fig 7.**
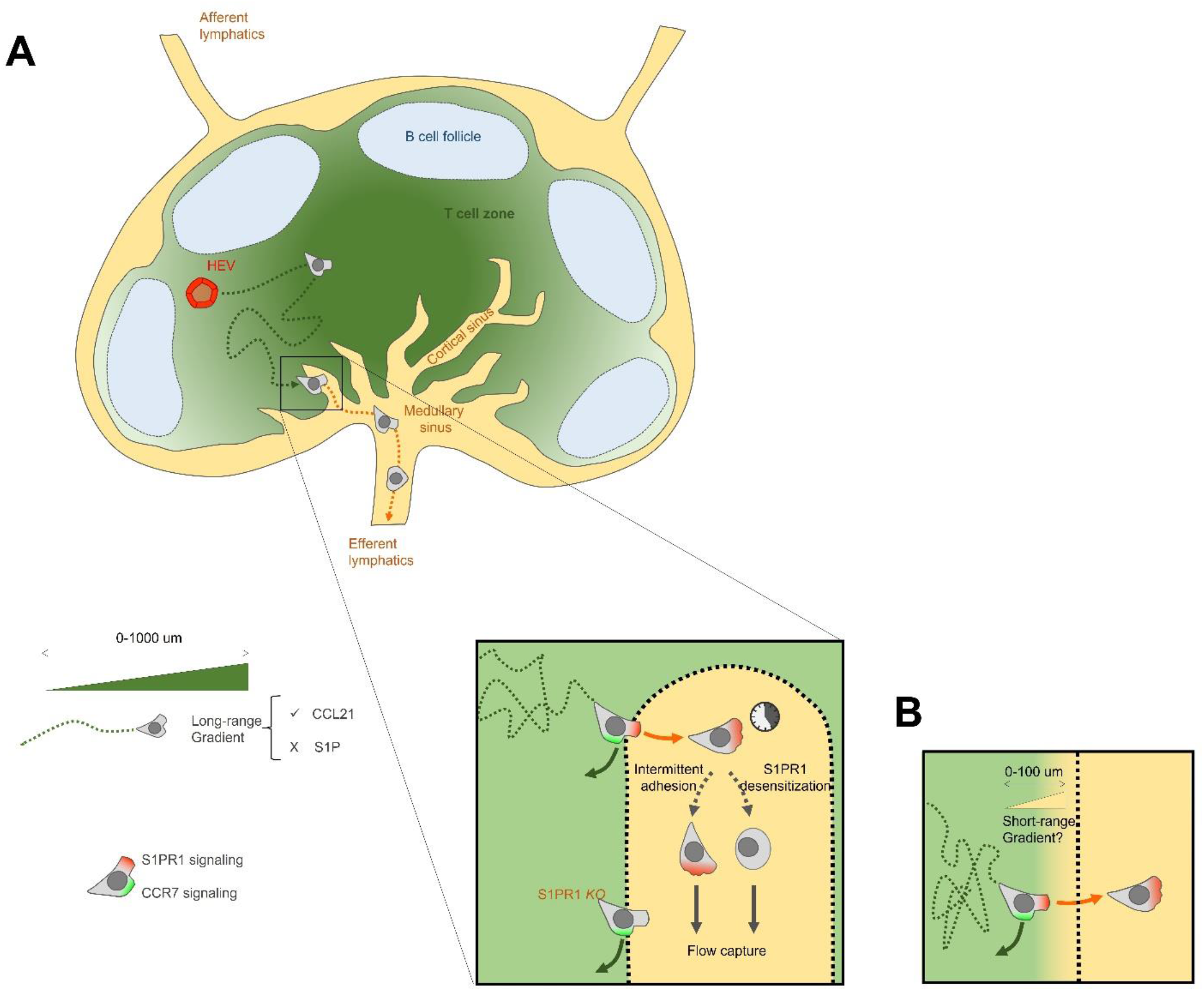
Minimal mechanistic model for naïve T lymphocyte traffic in lymph nodes. (A) Naïve T cells exit HEVs and migrate towards the central parenchyma guided by a long-range chemotaxis towards CCL21 (green). In the insert: Upon randomly encountering a cortical sinus, instant S1P sensing triggers the transmigration decision. Instead, S1PR1 *knock out* cells are retained in the parenchyma due to counteracting CCR7 signaling. Once in sinus, intermittent adhesion and S1PR1 desensitization can independently trigger flow-capture and lymph node exit through efferent lymphatics. (B) The alternative hypothesis of a short-range S1P gradient diffusing into the parenchyma barely changes the scenario. The position of the decision making is shifted from the lymphatic endothelium to a neighboring zone, where S1P is sensed.

**Supplementary Fig 1.**
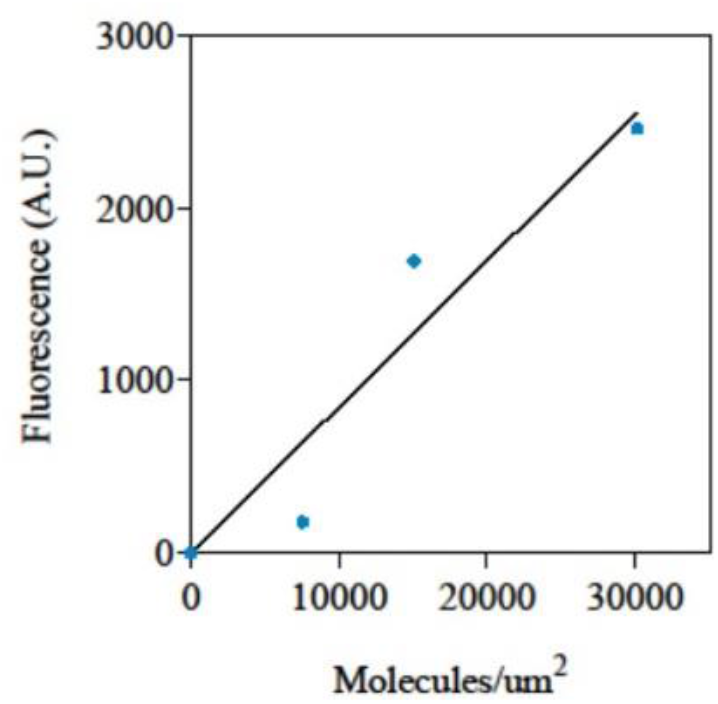
CCL21S6-Dy549P1 fluorescence intensity, measured in the same optic conditions as in Fig 3E and 3F, as a function of its surface concentration.

**Supplementary Fig 2.**
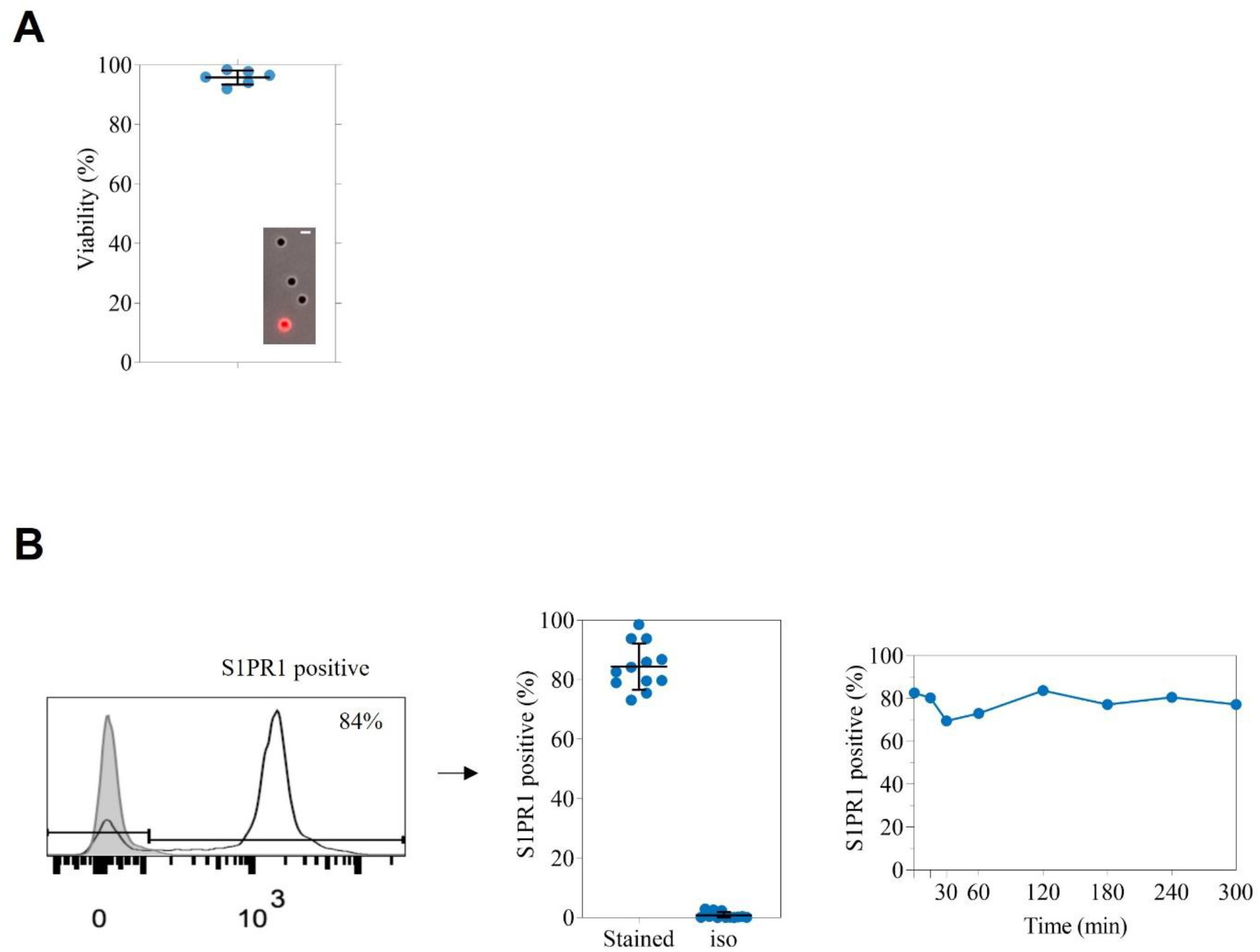
(A) Viability of cells from 6 independently tested donors after >6hs incubation without serum. Inset presents the live and dead conditions, scored by propidium iodide uptake (red cell). Scalebar = 10μm (B) Left and center, purified naïve T lymphocytes from 12 independently tested donors were fixed, stained for S1PR1, and the percent of S1PR1 positive cells was scored. The gray histogram corresponds to the isotype control used to define the gates. Right, S1PR1 expression vs time, for cells purified from 1 donor and incubated in the absence of serum for the indicated timepoints, then fixed and stained for S1PR1.

**Supplementary Fig 3.**
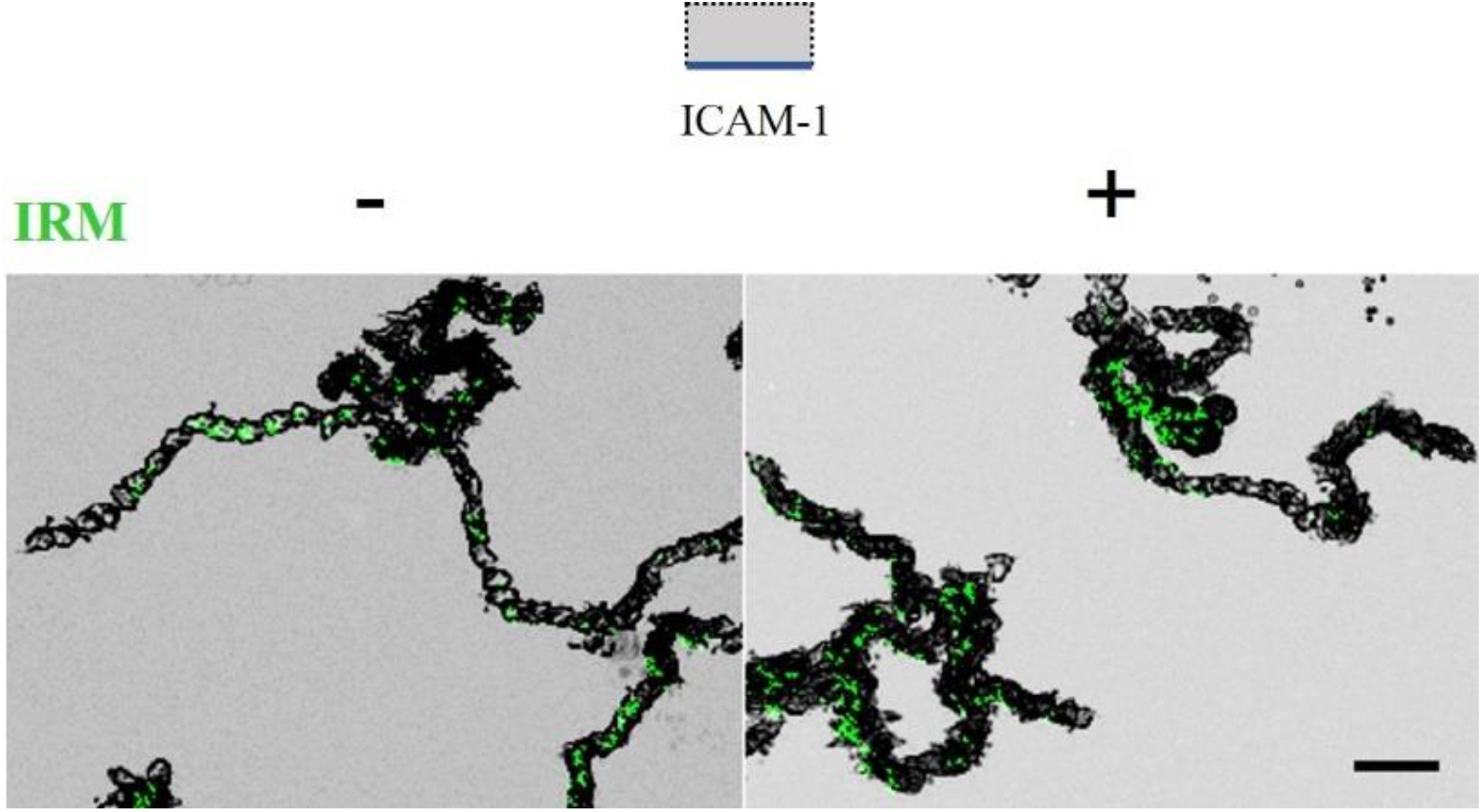
Overlaid bright field and IRM projections for cells migrating in the presence of 1μM CCL21 alone (-) or with 10% serum (+). Scalebar = 20μm.

## Discussion

Naïve T lymphocyte recirculation through lymph nodes is a critical feature of the immune system at steady state condition. It is required for the vast repertoire of cells to screen for a cognate antigen-loaded dendritic cell, and it represents a crucial therapeutic target for various pathologies. CCL21 and S1P orchestrate this continuous trafficking, as deduced from perturbation experiments and *in vivo* imaging, and have been claimed chemotactic. However, to our knowledge no migration assay along a single and controlled gradient, coupled to *live* imaging, has been reported so far for naïve T lymphocytes. Here, we used customized *in vitro* setups to perform *live* imaging of human naïve T lymphocytes migrating under defined environmental conditions. By exposing cells to controlled and single stimuli, we revisited several dogmas in the field of lymphocyte migration and provided a quantitative directionality analysis for migrating naïve T lymphocytes.

We first demonstrated that, contrary to the literature^14^, CCL21 signals on naïve T lymphocytes not only when adsorbed but also in bulk solution. Because chemokine adsorption is based on electrostatic interactions, our assays do not discriminate whether those molecules are truly read from the substrate or slowly desorbing into a local soluble pool, in which case the substrate would only act as a chemokine depot. Future assays with a molecule faithfully bound to the substrate, such as through a biotinylated linker, should help answering this question. Regardless the mechanism under play, we verified the widely accepted haptokinesis and haptotaxis of naïve T lymphocytes in response to adsorbed CCL21, and we recorded their chemokinesis and chemotaxis in response to bulk CCL21, providing fine directionality analysis for both cases of taxis. Chemotactic indexes were higher for soluble gradients than adsorbed ones, despite the narrower range of experienced slopes in the first case, indicating that soluble gradients may be more efficient at guiding cells. More generally, cell directivity proved in a similar range as that previously measured in response to CCL19^19^, but lower than that reported for dendritic cells migrating towards both CCR7 ligands^53,54^. Such values may reflect the importance of CCR7 guidance for antigen-loaded dendritic cells, for which a ballistic motion is needed to efficiently reach lymph nodes, whereas naïve T lymphocytes are gently attracted to areas of antigen presentation while keeping a screening behavior.

Our data also dismiss the reported lack of naïve T lymphocyte adhesion on ICAM-1^14^. By using IRM imaging, we proved that the speed of naïve T lymphocytes is modulated by intermittently adhering on ICAM-1. This is in line with *in vivo* data showing that LFA-1 is necessary to sustain fast migration in lymph nodes^34,55^. Interestingly, speed fluctuations were reported for migrating T lymphocytes in mouse lymph nodes^15^, and migration arrests were further correlated to intracellular calcium peaks *in vivo* and *in vitro*, at least for effector T lymphocytes in confined environments^56^. Given that human neutrophils also modulate their adhesion while migrating on ICAM-1 substrates^44^, it is tempting to speculate whether both cell types bear an internal clock controlling such adhesion runs, hence representing an intrinsic and intended feature. In the case of naïve T lymphocytes, perhaps to avoid excessive adhesion on ICAM-1-expressing dendritic cells and ensure antigen screening continuation. Simultaneous IRM and calcium imaging during *in vitro* migration should help answering this question. Finally, intermittent adhesion was not stabilized by an excess of bulk chemokine nor mild flow. A hypothesis is that stable adhesion under flow may be achieved by adding a selectin-mediated rolling step, in line with the long-established adhesion cascade for leukocyte transmigration through High Endothelial Venules (HEVs)^57^.

Finally, we analyzed the real-time response of naïve T lymphocytes to S1P-rich serum. After almost two decades or research, S1P biology remains a controversial topic with several standing models explaining lymphocyte exit from lymph nodes. Models claiming chemotaxis towards S1P are based on weak *in vitro* transmigration values, either due to premature exposure to suboptimal S1P concentrations and concomitant receptor desensitization, or lack of an appropriate carrier molecule. Here, by visualizing in real-time the cells’ reaction to gradients of bioactive S1P, we observed two different behaviors: a small fraction of cells transiently polarizing and migrating, but with a short displacement and only a mild skew in directionality towards the serum source, and a bigger fraction of cells transiently polarizing on the spot and without displacement. Altogether, these data do not support the existence of efficient S1P-triggered chemotaxis, since under the same conditions in which CCL21 gradients triggered durable and long-range chemotaxis, serum factors, hence S1P, did not. Interestingly, because our gradients are built by gradually diffusing factors and likely also causing premature S1PR1 desensitization, these behaviors matched the reported values from Transwell assays: the observed 8% migrating cells recapitulate the transmigration values reported towards S1P alone^4,30–34^, while the total 21% polarizing cells recapitulate the transmigration values reported towards S1P together with lymphatic endothelial cells (LECs)^35^. This correspondence suggested that cells polarizing in the spot represented a commitment to transmigrate through (and most likely aided by) LECs, and gave us the indication for an optimal S1P-sensing condition. Indeed, when cells were instantly exposed to serum the percent of polarizing cells increased to ~ 60%, to our knowledge the highest effect reported to date. Since migrating cells *in vivo* encounter cortical sinuses in an already polarized fashion, likely by CCR7 ligands, and since a similar effect was observed for 10% and 100% serum concentrations, we conclude serum factors provide naïve T lymphocytes with a qualitative ‘decision making’ signal, to trigger transmigration into cortical sinuses.

Taken together, our data complements *in vivo* observations and leads us to the following model of naïve T lymphocyte traffic (Fig. 7A). After accessing lymph nodes through HEVs or the subcapsular sinus floor, cells are gently attracted by long-range CCL19 and CCL21 gradients towards the central parenchyma. During this journey, they scan antigen-loaded dendritic cells, until randomly encountering a cortical sinus. Upon probing its lumen, S1PR1 is needed to achieve successful transmigration^15^. Since we did not observe any apparent serum-based inhibition of cell adhesion nor CCL21-triggered migration, the main role of S1P may be to force the transmigration step upon probing the cortical sinus lumen. Once in the lumen, intermittent adhesion and S1PR1 desensitization (followed by cell depolarization) can independently trigger the detachment of cells from LECs and their concomitant capture by the increasing flow of efferent lymph, which finally brings cells out of the lymph node and back into circulation. Our data does not exclude the possibility of a short-range S1P guidance (an order of magnitude lower than that of CCL21) in the vicinity of cortical sinuses (Fig. 7B). In such scenario the decision to exit may be taken upon gradient detection, a few body-lengths before reaching LECs, but the subsequent steps would remain the same. However, lack of *in vivo* evidence supporting a S1P gradient or chemotaxis towards sinuses, and rapid S1PR1 desensitization *in vitro* upon gradient sensing, often without displacement (Fig. 5), leads us to favor the first model. Finally, a potential stromal gate function of LECs is also not excluded, for instance by providing a scaffold for adhesion and transmigration or by delivering additional soluble or contact signals. Such notion is supported by the different, S1P-independent basal speed observed for cells migrating in the medullary cords as compared to those in the parenchyma^33^.

To conclude, *live* imaging in reductionist *in vitro* setups gave us a mechanistic understanding of naïve T lymphocyte traffic, complementing *in vivo* observations. Such insights may foster the development of more specific, tailored immunosuppressive drugs. Our study also paves the way for a consistent and quantitative characterization of human leukocyte migration. With a recent report on human B lymphocyte migration revealing differences with the mouse system^42^, this quest may become more relevant than previously thought.

## Acknowledgments and funding

We are grateful to the Cell Culture Platform facility (Luminy TPR2-INSERM), the Turing Centre for Living systems, the Regional Council of Provence-Alpes-Côte d’Azur and the LABEX INFORM program. Also the European Union’s Horizon 2020 research and innovation program under the Marie Skłodowska-Curie grant agreement No713750; Agence Nationale de la Recherche, RECRUTE - ANR-15-CE15-0022 and ILIAAD ANR-18-CE09-0029; A*MIDEX n° ANR-11-IDEX-0001-02; and The Swiss National Science Foundation, grant number 310030_189144.

## Author contributions

NGS and OT designed the experiments, SS performed haptotaxis experiments, LDB and NGS performed serum-triggered polarization experiments, NGS performed all remaining experiments, NGS and SS wrote the scripts for analysis and analyzed the data, CM and MA produced the recombinant CCL21 variants under DFL’s supervision, NGS wrote the manuscript, all remaining authors revised it. NGS and OT supervised the project.

## Ethics statement

Human subjects: Blood from healthy volunteers was obtained through a formalized agreement with the French Blood Agency (Etablissement Français du Sang, agreement n° 2017-7222). Blood was obtained by the agency after informed consent of the donors, in accordance with the Declaration of Helsinki. All experiments were approved by the INSERM Institutional Review Board and ethics committee.

## Materials and Methods

### Cells

Whole blood from healthy adult donors of group 0, drawn into EDTA tubes, was obtained from the “Établissement Français du Sang”. Peripheral Blood Mononuclear Cells (PBMC) were recovered from the interface of a Ficoll gradient (Eurobio, Evry, France) and washed with Phosphate Buffer Saline (PBS, Gibco). Naïve T lymphocytes were purified with the Miltenyi naïve CD4+ T Cell Isolation Kit II. After purification, cells were kept in RPMI 1640 medium supplemented with penicillin 100 U/ml (Gibco, Carlsbad, CA), streptomycin 100 μg/ml (Gibco, Carlsbad, CA), 25 mM GlutaMax (Gibco, Carlsbad, CA), with or without 10% fetal calf serum (FCS, Lonza, Basel, Switzerland) in a 37°C incubator with 5% CO_2_, until use. Human serum was prepared from the same donor, from blood coagulated in a dry tube. After 5 minutes centrifugation at 500 RCF the supernatant was taken, filtered through a 0.2 μm mesh and kept at 37°C until use.

### Flow cytometry

One hundred thousand cells per condition were fixed for 10 minutes with 1% paraformaldehyde (PFA, Thermofisher), then washed once with 4 mL of FACS buffer (2% FCS in PBS), resuspended in 100 μL and stained for 30 minutes at 4°C in the dark. They were finally washed with 4 mL FACS buffer and re-suspended in 0.5 mL to be analyzed in a LSR Fortessa X20 (BD Biosciences, Europe). For live staining, an equal number of purified cells or PBMCs were stained first on ice, then washed with FACS buffer and fixed with 1% PFA. For the blood sample, 200uL were first blocked with 1 mg human IgG (Tegeline, LFB Biomedicaments) for 15 minutes on ice, immediately after stained for 30 minutes, erythrocytes were then lysed with RBC lysis buffer (eBioscience), finally cells were washed with 12 mL PBS and fixed with 1% PFA. The antibodies used for staining were APC/Cyanine7 anti-human CD45RA (clone HI100, Biolegend), PE anti-human CD197 (CCR7, clone G043H7, Biolegend), PE/Cy7 anti-human CD62L (clone DREG-56, Biolegend), EF660 anti-human CD363 (S1PR1, clone SW4GYPP, eBioscience) and EF660 IGG1K isotype control (eBioscience).

### Devices

Single channel devices consisted of Ibidi μ-Slide uncoated IV 0.4 (Ibidi GMBH, Martinsreid, Germany). Surfaces were coated overnight at 4°C, either with 10 μg/mL human ICAM-1 (R&D Systems), 1 μg/mL CCL21 (Miltenyi biotech) or a mixture of both, followed by blocking with 4% fatty acid-free Bovine Serum Albumin (BSA, Sigma) solution in PBS, for at least 15 min at room temperature. Devices were finally rinsed with PBS and then culture media. Flow experiments were performed in single channels connected to a pump system (neMESYS 290N, Cetoni). The gradient device was molded in PDMS and prepared as described elsewhere^19^, 10 kDa Dextran FITC (Sigma) was used as a diffusion marker to analyze gradient dynamics. For CCL21 surface micropatterning, CCL21-S6^Dy549P1^ homogeneous substrates in single channels were UV-illuminated (λ= 375 nm, 300s exposition at 5V) through a Digital Micromirror Device (Primo; Alveole) in the presence of photoinitiator (PLPP, Alveole), to degrade the chemokine in a modulated way. Patterns of interest were designed on Matlab (The MathWorks). The fluorescent intensity was calibrated into number of molecules by imaging serial dilutions in 39 μm hight PDMS microchannels, passivated with PEG-SVA to limit adsorption, as described elsewhere^58^. Substrates for cell capture were prepared on glass slides (Schott High performance coverslip 22×22#1.5H cleanroom cleaned) which were plasma activated (Harrick Plasma) during 30min and then incubated for 2 hours at 4°C with a 1% APTES 0.03% Acetic acid solution. Slides were then rinsed with Milli-Q water, dried and baked 15 min at 95°C. Open-wells (6×2mm) were finally created by sticking a 250μm-thick PDMS sticker on the slides. A solution of 23% mPEG-SVA (MW:5000 Da, INTERCHIM) 10mM NaHCO_3_ was then added to the wells, and substrates were incubated overnight at 4°C. The next day they were rinsed with Milli-Q water, and 10μl of PLPP solution (Alveole) was added. Arrays of capture spots, 3μm in radius, were illuminated with UV (λ=375nm for 180sec, ~2488 mJ/mm^2^ dose) through a Digital Micromirror Device (Primo™, Alvéole). Wells were rinsed with PBS and incubated overnight with 50 μg/mL anti-human CD45RA antibody (Abcam ab212774, lot GR3365431-3), at 4°C.

### Recombinant chemokine expression

pSUMO ΔCCL21 was cloned using the primers 5’-GGTGCTCGAGTCAGCCCTGGGCTGGTTTCTGTGGGGATGGTGTCTTG-3’ and 5’-CCCTCTAGAAATAATTTTGTTTAACTTTAAGAAGGAGATATACATATGG-3’ to amplify ΔCCL21 from pSUMO CCL21^26^ and introducing it into the pSUMO backbone using XbaI and XhoI cutting sites. The chemokines and dyes were produced as previously described^46^. In brief, S6-tagged CCL21 and ΔCCL21 were each produced in E.coli BL21 DE3, refolded and using a multi step protocol affinity purified, with a final C18 reverse phase HPLC step. CoA-Dy549P1 was generated as described using DY-549P1-Maleimide (Dyomics GmbH, Germany) and CoA Li_3_ (Sigma-Aldrich, Switzerland). CoA-Dy549P1 was transferred to CCL21-S6 using the phosphopantetheinyl transferase Sfp and labelled chemokine purified using C18 reverse phase HPLC.

### Imaging and data analysis

Experiments were performed on an inverted Zeiss Z1 automated microscope (Carl Zeiss, Germany) equipped with a CoolSnap HQ CCD camera (Photometrics) and piloted by μManager 1.4. Plan-Apochromat 10x/0.3 and 20x/0.8 air objectives were used for bright-field imaging, while a 40x/1.3 oil one was used for Interference Reflection Microscopy (IRM). IRM images were first corrected by subtracting a background image, secondly the pixel values were inverted to convert dark signal into positive values, finally a rolling ball algorithm was applied to flatten the image. Cells were tracked using the FIJI plugin Trackmate^59^, except for serum experiments where the Manual Tracking plugin was used. Tracks were exported and further analysis and plots were performed with MATLAB custom-made scripts (MATLAB software, The MathWorks, Natick, MA, USA). The unbiased colormap for the heatmaps was taken from ref^60^. Gradient dynamics were analyzed with a custom-made FIJI macro, fluorescence intensity values were normalized with the equation:

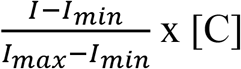

Where *I_min_* is the average value recorded on the sink channel (background), *I_max_* is the average value recorded on the source channel, and [C] is the chemokine concentration applied at the source. Ilastik^61^ was used for morphometric analysis, to create binary masks which were further processed with Fiji and analyzed with Matlab custom scripts.

### Statistical Analysis

Multiple comparison ANOVA tests were used to compare datasets. The number of analyzed cells, tested donors and resulting *p*-values are specified on each figure’s legend.

## References

1. von Andrian, U. H. & Mempel, T. R. Homing and cellular traffic in lymph nodes. Nat Rev Immunol 3, 867–878 (2003).

2. Schulz, O., Hammerschmidt, S. I., Moschovakis, G. L. & Förster, R. Chemokines and Chemokine Receptors in Lymphoid Tissue Dynamics. Annu. Rev. Immunol. 34, 203–242 (2016).

3. Baeyens, A. A. L. & Schwab, S. R. Finding a Way Out: S1P Signaling and Immune Cell Migration. Annu. Rev. Immunol. 38, 759–784 (2020).

4. Pham, T. H. M., Okada, T., Matloubian, M., Lo, C. G. & Cyster, J. G. S1P1 Receptor Signaling Overrides Retention Mediated by Gαi-Coupled Receptors to Promote T Cell Egress. Immunity 28, 122–133 (2008).

5. Tsai, H.-C. & Han, M. H. Sphingosine-1-Phosphate (S1P) and S1P Signaling Pathway: Therapeutic Targets in Autoimmunity and Inflammation. Drugs 76, 1067–1079 (2016).

6. Maceyka, M., Harikumar, K. B., Milstien, S. & Spiegel, S. Sphingosine-1-phosphate signaling and its role in disease. Trends in Cell Biology 22, 50–60 (2012).

7. Okada, T. et al. Antigen-Engaged B Cells Undergo Chemotaxis toward the T Zone and Form Motile Conjugates with Helper T Cells. PLoS Biology 3, 15 (2005).

8. Ulvmar, M. H. et al. The atypical chemokine receptor CCRL1 shapes functional CCL21 gradients in lymph nodes. Nat Immunol 15, 623–630 (2014).

9. Braun, A. et al. Afferent lymph–derived T cells and DCs use different chemokine receptor CCR7–dependent routes for entry into the lymph node and intranodal migration. Nat Immunol 12, 879–887 (2011).

10. Rot, A. & von Andrian, U. H. Chemokines in innate and adaptive host defense: basic chemokinese grammar for immune cells. Annu Rev Immunol 22, 891–928 (2004).

11. Yang, B.-G. et al. Binding of Lymphoid Chemokines to Collagen IV That Accumulates in the Basal Lamina of High Endothelial Venules: Its Implications in Lymphocyte Trafficking. J Immunol 179, 4376–4382 (2007).

12. de Paz, J. L. et al. Profiling Heparin–Chemokine Interactions Using Synthetic Tools. ACS Chem. Biol. 2, 735–744 (2007).

13. Weber, M. et al. Interstitial Dendritic Cell Guidance by Haptotactic Chemokine Gradients. Science 339, 328–332 (2013).

14. Woolf, E. et al. Lymph node chemokines promote sustained T lymphocyte motility without triggering stable integrin adhesiveness in the absence of shear forces. Nat Immunol 8, 1076–1085 (2007).

15. Miller, M. J., Wei, S. H., Cahalan, M. D. & Parker, I. Autonomous T cell trafficking examined in vivo with intravital two-photon microscopy. Proceedings of the National Academy of Sciences 100, 2604–2609 (2003).

16. Grigorova, I. L. et al. Cortical sinus probing, S1P 1-dependent entry and flow-based capture of egressing T cells. Nature Immunology 10, 58–65 (2009).

17. Okada, T. & Cyster, J. G. CC Chemokine Receptor 7 Contributes to Gi-Dependent T Cell Motility in the Lymph Node. The Journal of Immunology 178, 2973–2978 (2007).

18. Worbs, T., Mempel, T. R., Bölter, J., von Andrian, U. H. & Förster, R. CCR7 ligands stimulate the intranodal motility of T lymphocytes in vivo. The Journal of Experimental Medicine 204, 489–495 (2007).

19. Garcia-Seyda, N. et al. Microfluidic device to study flow-free chemotaxis of swimming cells. Lab Chip (2020) doi:10.1039/D0LC00045K.

20. Jørgensen, A. S. et al. CCL19 with CCL21-tail displays enhanced glycosaminoglycan binding with retained chemotactic potency in dendritic cells. J Leukoc Biol 104, 401–411 (2018).

21. Luther, S. A., Tang, H. L., Hyman, P. L., Farr, A. G. & Cyster, J. G. Coexpression of the chemokines ELC and SLC by T zone stromal cells and deletion of the ELC gene in the plt/plt mouse. Proceedings of the National Academy of Sciences 97, 12694–12699 (2000).

22. Link, A. et al. Fibroblastic reticular cells in lymph nodes regulate the homeostasis of naive T cells. Nat Immunol 8, 1255–1265 (2007).

23. Miller, H. et al. High-Speed Single-Molecule Tracking of CXCL13 in the B-Follicle. Front. Immunol. 9, (2018).

24. Cosgrove, J. et al. B cell zone reticular cell microenvironments shape CXCL13 gradient formation. Nat Commun 11, 3677 (2020).

25. Schumann, K. et al. Immobilized Chemokine Fields and Soluble Chemokine Gradients Cooperatively Shape Migration Patterns of Dendritic Cells. Immunity 32, 703–713 (2010).

26. Hauser, M. A. et al. Distinct CCR7 glycosylation pattern shapes receptor signaling and endocytosis to modulate chemotactic responses. Journal of Leukocyte Biology 99, 993–1007 (2016).

27. Graham, G. J., Handel, T. M. & Proudfoot, A. E. I. Leukocyte Adhesion: Reconceptualizing Chemokine Presentation by Glycosaminoglycans. Trends in Immunology 40, 472–481 (2019).

28. Ramos-Perez, W. D., Fang, V., Escalante-Alcalde, D., Cammer, M. & Schwab, S. R. A map of the distribution of sphingosine 1-phosphate in the spleen. Nat Immunol 16, 1245–1252 (2015).

29. Fang, V. et al. Gradients of the signaling lipid S1P in lymph nodes position natural killer cells and regulate their interferon-γ response. Nat Immunol 18, 15–25 (2017).

30. Arnon, T. I. et al. GRK2-Dependent S1PR1 Desensitization Is Required for Lymphocytes to Overcome Their Attraction to Blood. Science 333, 1898–1903 (2011).

31. Matloubian, M. et al. Lymphocyte egress from thymus and peripheral lymphoid organs is dependent on S1P receptor 1. Nature 427, 355–360 (2004).

32. Schwab, S. R. Lymphocyte Sequestration Through S1P Lyase Inhibition and Disruption of S1P Gradients. Science 309, 1735–1739 (2005).

33. Drouillard, A. et al. Human Naive and Memory T Cells Display Opposite Migratory Responses to Sphingosine-1 Phosphate. J.I. 200, 551–557 (2018).

34. Reichardt, P. et al. A role for LFA-1 in delaying T-lymphocyte egress from lymph nodes. EMBO J 32, 829–843 (2013).

35. Xiong, Y. et al. CD4 T cell sphingosine 1-phosphate receptor (S1PR)1 and S1PR4 and endothelial S1PR2 regulate afferent lymphatic migration. Sci. Immunol. 4, eaav1263 (2019).

36. Graeler, M., Shankar, G. & Goetzl, E. J. Cutting Edge: Suppression of T Cell Chemotaxis by Sphingosine 1-Phosphate. J Immunol 169, 4084–4087 (2002).

37. Ledgerwood, L. G. et al. The sphingosine 1-phosphate receptor 1 causes tissue retention by inhibiting the entry of peripheral tissue T lymphocytes into afferent lymphatics. Nat Immunol 9, 42–53 (2008).

38. Wei, S. H. et al. Sphingosine 1-phosphate type 1 receptor agonism inhibits transendothelial migration of medullary T cells to lymphatic sinuses. Nat Immunol 6, 1228–1235 (2005).

39. Aoun, L. et al. Leukocyte transmigration and longitudinal forward-thrusting force in a microfluidic Transwell device. Biophys J 120, 2205–2221 (2021).

40. Davis, M. M. A Prescription for Human Immunology. Immunity 29, 835–838 (2008).

41. Pulendran, B. & Davis, M. M. The science and medicine of human immunology. 13 (2020).

42. Park, S. M. et al. Migratory cues controlling B-lymphocyte trafficking in human lymph nodes. Immunology & Cell Biology 99, 49–64 (2021).

43. Aoun, L. et al. Amoeboid Swimming Is Propelled by Molecular Paddling in Lymphocytes. Biophysical Journal (2020) doi:10.1016/j.bpj.2020.07.033.

44. Garcia-Seyda, N. et al. Human neutrophils swim and phagocytise bacteria. Biology of the Cell 113, 28–38 (2021).

45. Barry, N. P. & Bretscher, M. S. Dictyostelium amoebae and neutrophils can swim. Proceedings of the National Academy of Sciences 107, 11376–11380 (2010).

46. Artinger, M., Matti, C., Gerken, O. J., Veldkamp, C. T. & Legler, D. F. A Versatile Toolkit for Semi-Automated Production of Fluorescent Chemokines to Study CCR7 Expression and Functions. International Journal of Molecular Sciences 22, 4158 (2021).

47. Luo, X. et al. Lymphocytes perform reverse adhesive haptotaxis mediated by LFA-1 integrins. J Cell Sci 133, (2020).

48. Strale, P.-O. et al. Multiprotein Printing by Light-Induced Molecular Adsorption. Adv. Mater. 28, 2024–2029 (2016).

49. Christoffersen, C. et al. Endothelium-protective sphingosine-1-phosphate provided by HDL-associated apolipoprotein M. PNAS 108, 9613–9618 (2011).

50. Blaho, V. A. et al. HDL-bound sphingosine-1-phosphate restrains lymphopoiesis and neuroinflammation. Nature 523, 342–346 (2015).

51. Schwab, S. R. et al. Lymphocyte Sequestration Through S1P Lyase Inhibition and Disruption of S1P Gradients. Science 309, 1735–1739 (2005).

52. Lo, C. G., Xu, Y., Proia, R. L. & Cyster, J. G. Cyclical modulation of sphingosine-1-phosphate receptor 1 surface expression during lymphocyte recirculation and relationship to lymphoid organ transit. Journal of Experimental Medicine 201, 291–301 (2005).

53. Aizel, K. et al. A tuneable microfluidic system for long duration chemotaxis experiments in a 3D collagen matrix. Lab Chip 17, 3851–3861 (2017).

54. Vargas, P. et al. Innate control of actin nucleation determines two distinct migration behaviours in dendritic cells. Nat Cell Biol 18, 43–53 (2016).

55. Katakai, T., Habiro, K. & Kinashi, T. Dendritic Cells Regulate High-Speed Interstitial T Cell Migration in the Lymph Node via LFA-1/ICAM-1. J.I. 191, 1188–1199 (2013).

56. Dong, T. X. et al. Intermittent Ca2+ signals mediated by Orai1 regulate basal T cell motility. eLife 6, e27827 (2017).

57. Zhou, F. et al. The kinetics of E-selectin- and P-selectin-induced intermediate activation of integrin αLβ2 on neutrophils. Journal of Cell Science 134, jcs258046 (2021).

58. Robert, P. et al. Functional Mapping of Adhesiveness on Live Cells Reveals How Guidance Phenotypes Can Emerge From Complex Spatiotemporal Integrin Regulation. Front. Bioeng. Biotechnol. 9, 625366 (2021).

59. Tinevez, J.-Y. et al. TrackMate: An open and extensible platform for single-particle tracking. Methods 115, 80–90 (2017).

60. Crameri, F., Shephard, G. E. & Heron, P. J. The misuse of colour in science communication. Nature Communications 11, 5444 (2020).

61. Berg, S. et al. ilastik: interactive machine learning for (bio)image analysis. Nat Methods 16, 1226–1232 (2019).

